# Complex interactions between local adaptation, plasticity, and sex affect vulnerability to warming in a widespread marine copepod

**DOI:** 10.1101/423624

**Authors:** Matthew Sasaki, Sydney Hedberg, Kailin Richardson, Hans G. Dam

**Affiliations:** Department of Marine Science, University of Connecticut, Groton, CT, USA; Gustavus Adolphus College, St. Peter, MN, 56082; Savannah State University, Savannah, GA, 31404

**Keywords:** Thermal adaptation, climate change, copepod, sex-specific response, developmental plasticity, Acartia tonsa

## Abstract

Predicting the response of populations to climate change requires knowledge of thermal performance. Genetic differentiation and phenotypic plasticity affect thermal performance, but the effects of sex and developmental temperatures often go uncharacterized. We used common garden experiments to test for effects of local adaptation, developmental phenotypic plasticity, and individual sex on thermal performance of the ubiquitous copepod, *Acartia tonsa*. Females had higher thermal tolerance than males in both populations, while the Florida population had higher thermal tolerance compared to the Connecticut population. An effect of developmental phenotypic plasticity on thermal tolerance was observed only in the Connecticut population. Ignoring sex-specific differences may result in a severe underestimation of population-level impacts of warming (i.e. - population decline due to sperm limitation). Further, despite having a higher thermal tolerance, southern populations may be more vulnerable to warming as they lack the ability to respond to increases in temperature through phenotypic plasticity.

## Introduction

Temperature has a profound effect on organismal performance [1,2]. Rapid climate warming represents a significant challenge for organisms, increasing average environmental temperatures [3] and the frequency of extreme climatic events (i.e. - heat waves) [4]. Predicting organismal responses to these changes depends on our understanding of the factors affecting thermal tolerance. Acute thermal tolerance is known to be affected by phenotypic plasticity [5] and genetic differentiation [6], as well as diet, behavior and individual sex [7,8,9]. Spatial variation in the thermal environment should generate adaptive differences in thermal performance between populations from different environments [2]. The Climate Variability Hypothesis (CVH) [10,11] states that increased thermal tolerance should correspond with increased mean environmental temperature, while plasticity should evolve in response to variability in the thermal environment. This is supported in terrestrial and freshwater aquatic systems [12, 13], but there have been fewer tests in marine systems [14,15,16], yielding only limited support. Additionally, no tests of the CVH have examined how individual sex factors into observed variation in thermal performance.

Here we examine the effects of genetic differentiation, developmental phenotypic plasticity, and individual sex on thermal tolerance in the widespread copepod *Acartia tonsa*. This species dominates coastal and estuarine systems around the globe. With a geographic range covering a large latitudinal thermal gradient, this is a good model system to explore the contributions of various adaptive mechanisms to thermal adaptation [15]. Our results show that complex interactions between these variables strongly affect our ability to predict organismal responses to climate change.

## Methods

Plankton samples were collected from surface tows at field sites in Groton, Connecticut, and Punta Gorda, Florida (Table 1) during July and August of 2017 using a 250 um mesh plankton net. Temperature data for both sites was obtained from the AQUA-MODIS satellite database [17]. Initial laboratory populations, comprised of >1500 mature adults to minimize the effects of genetic drift, were established from collected animals. Cultures were maintained under common garden conditions (30 psu, 12:12 light:dark, 18°C) for several generations, and fed a diet of the microalgae *Tetraselmis* sp., *Rhodomonas* sp., and *Thalassiosira weissflogii*. Eggs were collected from the 18°C culture and moved to 22°C to develop. All other factors were held constant. Once mature, individuals from both developmental conditions (18 and 22°C) were exposed to a 24-hour acute heat stress. Individuals were carefully transferred to a microcentrifuge tube with 1.5 mL of filtered seawater, then transferred to heat blocks set to a constant temperature (18-38°C at degree intervals). Each individual experienced a single temperature. Individual survivorship was measured after 24 hours as binary data (1 = survival, 0 = mortality).

**Table 1.** – Site name, geographic coordinates, mean annual temperature, mean annual maximum, and mean annual temperature range for all collection locations.

| Population | Coordinates | Mean Annual<br>Temperature<br>(°C) | Mean<br>Maximum<br>Temperature<br>(°C) | Mean Annual<br>Temperature<br>Range (°C) |
| --- | --- | --- | --- | --- |
| Connecticut (CT) | 41.320591, -72.001564 | 13.3 | 22.7 | 22.5 |
| Florida (FL) | 26.940398, -82.051036 | 24.9 | 31.4 | 15.3 |

All analyses were performed using R version 3.5.1 [18]. 1717 individuals were used throughout the experiments. Initially, an ANOVA was run for all data (Survivorship ~ Stress Temperature + Sex + Developmental Temperature + Population, and all two-way interactions). Three-way and four-way interaction were excluded from the ANOVA. ANOVAs were also run for each population separately (Survivorship ~ Stress Temperature ^∗^ Sex ^∗^ Developmental Temperature). Thermal performance curves were estimated using logistic regressions. Because of the common garden design, differences in the performance curves between developmental conditions within a population can be attributed to developmental phenotypic plasticity, whereas differences between populations should reflect the effects of genetic differentiation. LD_50_ (the temperature with 50% mortality) was calculated for each performance curve. The change in LD50 between the two developmental conditions (ΔLD_50_) was used as a measure of the magnitude of the plastic response.

## Results

Statistical analyses – In the full ANOVA, significant effects of stress temperature, sex, developmental temperature and population, as well as significant stress temperature x population and developmental temperature x population interaction terms were evident (Table 2). Sex, stress temperature and developmental temperatures were significant in the Connecticut population, with no significant interaction terms. By contrast, only stress temperature and sex and their interaction were significant in the Florida population.

**Table 2.** - ANOVA results for the logistic regression relating survivorship to stress temperature, population, developmental temperature, and individual sex. Significant terms (p<0.05) are bolded.

|  | Df | Deviance | Resid. Df | Resid. Dev | Pr(>Chi) |
| --- | --- | --- | --- | --- | --- |
| **Stress Temperature** | 1 | 928.89 | 1715 | 1409.7 | 0.00 |
| **Individual Sex** | 1 | 54.7 | 1714 | 1355 | 1.40E-13 |
| **Developmental Temperature** | 1 | 7.74 | 1713 | 1347.2 | 0.005388 |
| **Population** | 1 | 17.8 | 1712 | 1329.4 | 2.45E-05 |
| **Dev. Temp. x Population** | 1 | 5.46 | 1711 | 1324 | 0.019462 |
| Sex x Dev. Temp | 1 | 0.27 | 1710 | 1323.7 | 0.60482 |
| Sex x Population | 1 | 3.82 | 1709 | 1319.9 | 0.050578 |
| **Stress Temp. x Population** | 1 | 8.38 | 1708 | 1311.5 | 0.003801 |
| Stress Temp. x Dev. Temp. | 1 | 0.06 | 1707 | 1311.5 | 0.809781 |
| Stress Temp. x Sex | 1 | 2.29 | 1706 | 1309.2 | 0.130522 |

CT Model
|  | Df | Deviance | Resid. Df | Resid. Dev | Pr(>Chi) |
| --- | --- | --- | --- | --- | --- |
| **Stress Temperature** | 1 | 318.6 | 725 | 666.22 | 0.00 |
| **Individual Sex** | 1 | 29.42 | 724 | 636.8 | 5.82E-08 |
| **Developmental Temperature** | 1 | 10.49 | 723 | 626.31 | 0.001202 |
| Stress Temp. x Sex | 1 | 0 | 722 | 626.31 | 0.960815 |
| Stress Temp. x Dev. Temp. | 1 | 0.14 | 721 | 626.17 | 0.71148 |
| Sex x Dev. Temp. | 1 | 0.23 | 720 | 625.94 | 0.631601 |
| Stress Temp. x Sex x Dev. Temp. | 1 | 1.75 | 719 | 624.2 | 0.186143 |

FL Model
|  | Df | Deviance | Resid. Df | Resid. Dev | Pr(>Chi) |
| --- | --- | --- | --- | --- | --- |
| **Stress Temperature** | 1 | 646.63 | 988 | 706.5 | 0.00 |
| **Individual Sex** | 1 | 20.51 | 987 | 685.99 | 5.94E-06 |
| Developmental Temperature | 1 | 0.63 | 986 | 685.36 | 0.42597 |
| **Stress Temp. x Sex** | 1 | 4.86 | 985 | 680.5 | 0.02744 |
| Stress Temp. x Dev. Temp. | 1 | 2.97 | 984 | 677.52 | 0.08465 |
| Sex x Dev. Temp. | 1 | 0.22 | 983 | 677.31 | 0.64267 |
| Stress Temp. x Sex x Dev. Temp. | 1 | 0.23 | 982 | 677.08 | 0.63395 |

Thermal performance curves - Males showed lower survival than females in both populations (Fig. 1, red versus blue lines). Developmental temperature had a strong effect on survival in the Connecticut population, but not in the Florida population.

**Figure 1.**
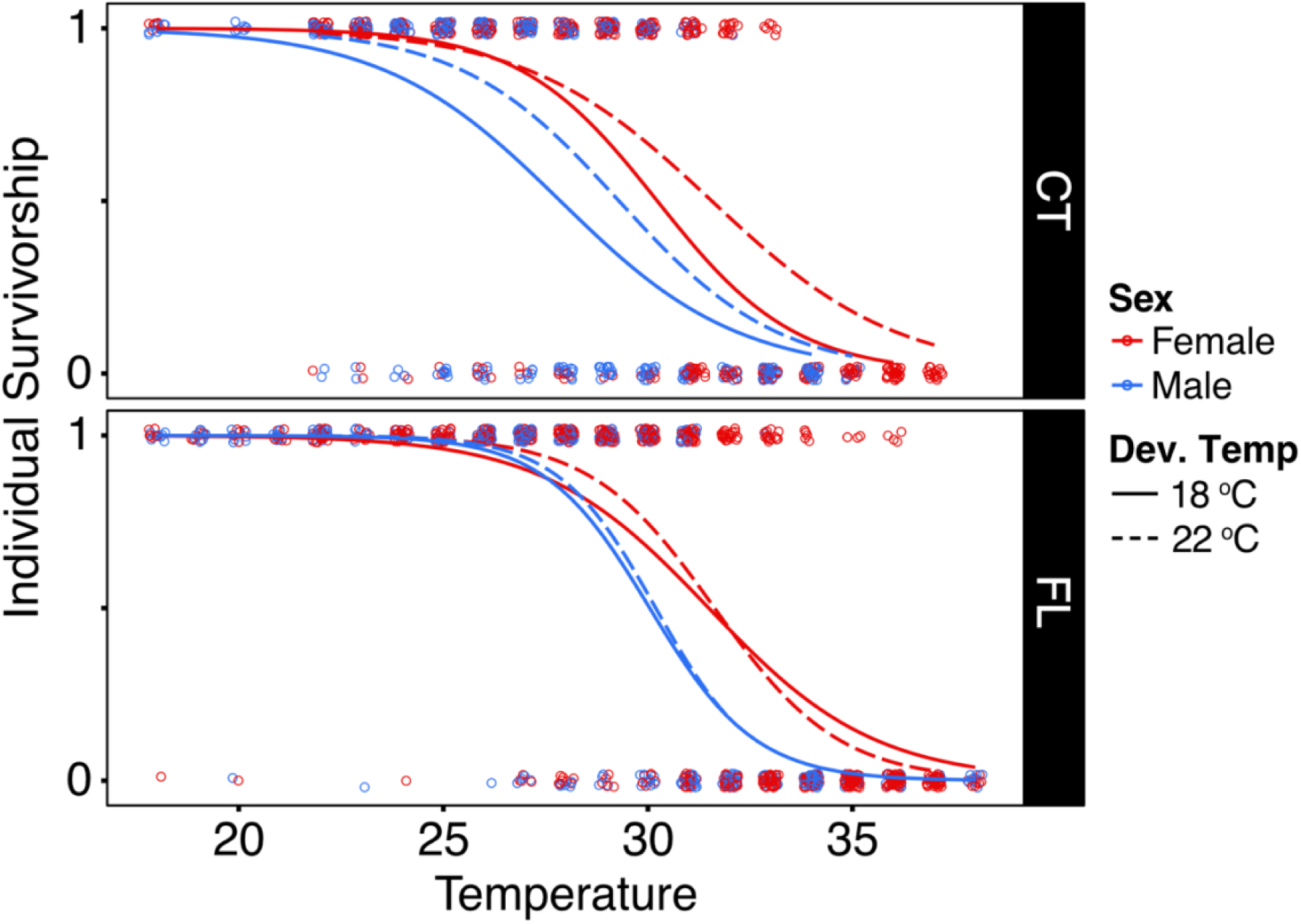
- Survivorship data for *Acartia tonsa* individuals is indicated by the points (1 = survival, 0 = mortality). Thermal performance curves are estimated by logistic regression. Color and line type indicate individual sex and developmental temperature respectively.

### Reaction norms –

Thermal tolerance (LD_50_) reaction norms show clear sex-dependent differences in thermal tolerance (Fig. 2), with females being always more tolerant than males. Higher developmental temperature increased thermal tolerance in the Connecticut population, but not in the Florida population. However, there were no sex-dependent differences in the plastic response between males and females (no difference in slopes), regardless of population.

**Figure 2.**
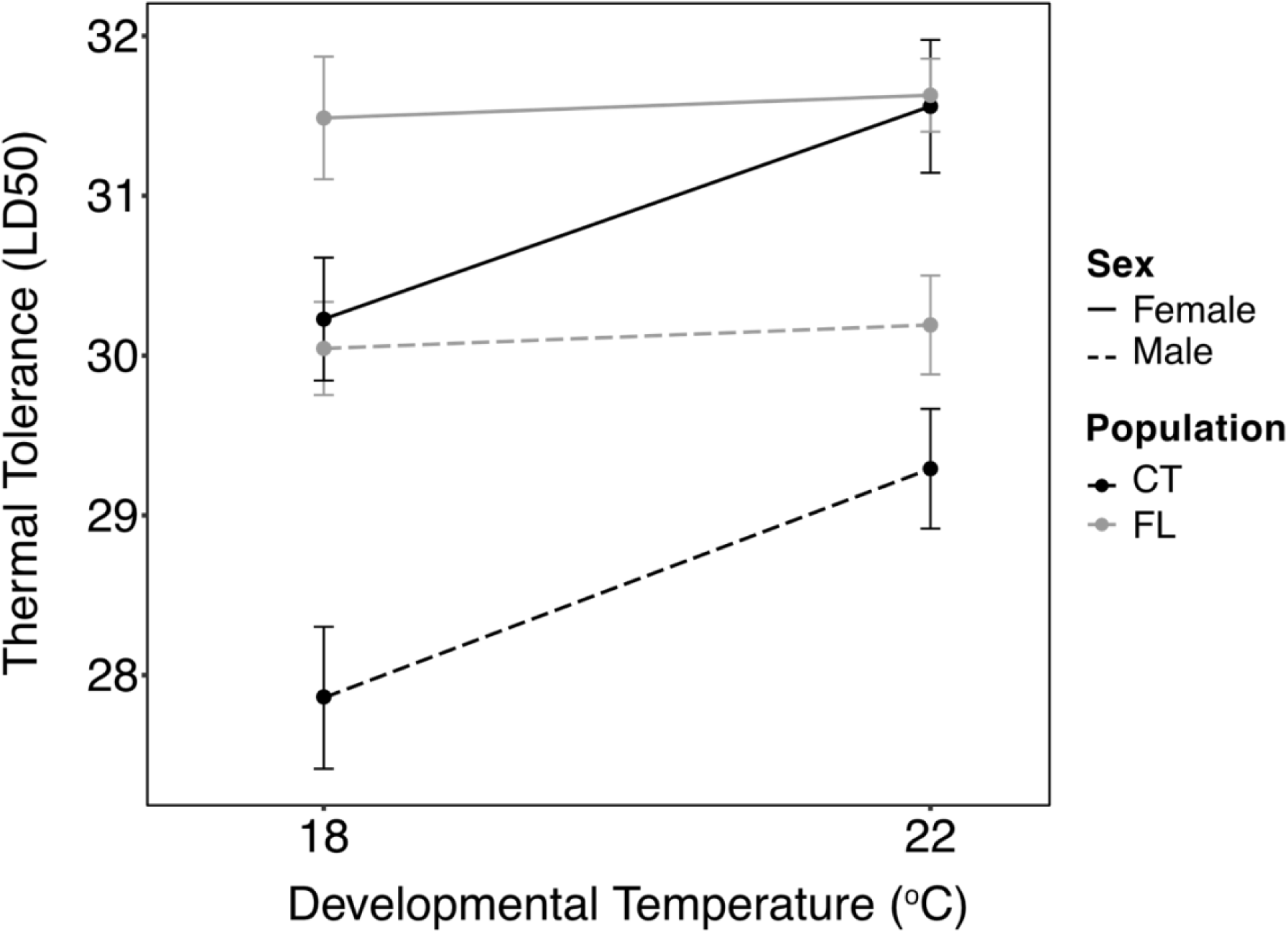
- Reaction norms showing thermal tolerance (LD_50_) as a function of developmental temperature for both sexes (line type) from the two populations (color). Points are thermal tolerance +/- SE from the logistic regression models. Reaction norm slope is the magnitude of plasticity.

## Discussion

The two populations of *Acartia tonsa* used in this study were collected from strongly differing thermal environments - a cool, variable environment (Connecticut), and a warmer, more stable environment (Florida). We observed lower thermal tolerance, but stronger plasticity in the Connecticut population relative to the Florida population, consistent with expectations of the CVH [10,11]. However, we find that individual sex had the largest effect on thermal tolerance. The demographic implications of these results are crucial to consider in predictions of future population dynamics.

In both populations, females always had a higher thermal tolerance than males. Sex-specific differences in thermal tolerance are observed across diverse systems [8, 19-21]. Within copepods, the few studies that have examined sex-specific thermal tolerance have also found females to be more thermally tolerant than males [9, 22-24], but ours is the first to examine these differences in more than one adaptive mechanism (thermal tolerance and phenotypic plasticity), and in multiple populations. Interestingly, *Acartia tonsa* females have also been found to be more tolerant to toxic dinoflagellates and to starvation [25, 26]. While there are strong differences in male and female thermal tolerance in this study, neither population exhibits significant differences between male and female plastic capacity (ΔLD_50_). No previous studies have examined differences in male and female developmental phenotypic plasticity, but higher acclimatory capacity was observed in females in a different copepod species [9]. This difference in sex-dependence of the different adaptive mechanisms suggests that their physiological basis is different.

Multiple factors affect acute thermal stress tolerance. Understanding these factors, and how they vary among populations, has critical implications for predictions of future population dynamics. Lower male thermal tolerance creates an asymmetrical vulnerability to climate change, which could lead to population declines under less intense warming due to sperm-limitation [27]. Our results also suggest that, despite having a higher thermal tolerance, southern populations may be more vulnerable to predicted warming. The southern population exists near their thermal limit under present conditions. As they are also unable to respond to increased temperatures through developmental phenotypic plasticity, any further increase in temperature is likely to have significant negative effects on population survival. Furthermore, males have a significantly lower thermal tolerance, lowering the temperature threshold that would bring the onset of temperature-driven population dynamic changes. Both sexes in the Connecticut population, however, have thermal tolerance values well above the current temperature maximum, and have a robust plastic capacity to increase thermal tolerance, potentially decreasing deleterious effects of warming on this population. Without examining multiple adaptive mechanisms and factors affecting thermal tolerance in several different populations, the complex variation in vulnerability to warming cannot be predicted.

## Acknowledgments

S.H. and K.R. were UCONN-Mystic Aquarium REU Program participants. We’d like to thank Drs. Tracy Romano and Michael Finiguerra for organizing this program.

## Author Contributions

M.S. collected and cultured all copepods. H.G.D. and M.S. designed the study. K.R. and S.H. performed the experiments. M.S. analyzed the data. M.S. and H.G. Dam drafted the manuscript.

## Data Accessibility

All data is accessible on Dryad (doi:10.5061/dryad.v5g6r80)

## Ethics Statement

Permits to collect individuals were not required. Standard protocols were used for copepod culturing and thermal tolerance assays. Ethics committee approval was not required.

